# Variable information across SNPs in GWAS data can cause false rejections of colocalisation which can be resolved by proportional colocalisation tests

**DOI:** 10.1101/2025.09.08.674910

**Authors:** Chris Wallace, Chloe Robins, Toby Johnson

## Abstract

Fine-mapping is now a standard post-GWAS analysis, but it has been shown to be potentially inaccurate for large meta-analysis GWAS. We show how this can be caused by variable amounts of statistical information between variants, e.g. due to variable sample sizes, which arise widely in meta-analyses, or due to variable imputation accuracy or call rate. This effect becomes more pronounced as sample sizes become larger, such that small differences in information available at different variants can lead to large errors in credible set predictions. Here, we show this inaccuracy propagates to the colocalisation method coloc, yielding confident but incorrect inference. We propose an adaptation of the proportional colocalisation test, which avoids any fine-mapping inference as an alternative to fine-mapping based colocalisation. This method was originally superseded by coloc because the null hypothesis is colocalization, and because it can have low power in genetic regions with many variants. Our adaptation resolves this second issue, and simulations demonstrate that our approach accurately resolves false coloc inferences. However, the first issue remains, such that we suggest proportional colocalisation is used as an accompaniment to fine-mapping-based coloc. Our findings also emphasize the need for new methodologies in fine-mapping, particularly as GWAS sample sizes grow, to ensure accurate fine-mapping inference.

## Introduction

Genome-wide association studies (GWAS) are designed to identify associations between genetic variation and human phenotypes. Post-GWAS analyses such as fine-mapping and colocalisation are conducted to move beyond association and identify the likely causal variants and genes underlying GWAS signals. Fine-mapping typically provides not only the posterior probability that any individual variant is causal, but also defines a credible set of variants, which are expected to contain the causal variant with some specified probability. Much stead is put in credible sets of size 1, as these are expected to identify with high probability the exact causal variant underlying a signal. Such findings open a path to experimental work to define the causal gene and mechanism based on comparing cellular or higher-level read-outs between different genotypes at that variant.

Fine-mapping methods typically involve calculating the evidence for a hypothesis of association versus no association at each variant or combination of variants, in the form of a Bayes factor (BF). These BF are then compared across variants to estimate the posterior probability of each variant being causal. The BF may be derived from single variant summary statistics (effect sizes and their standard errors)^1^ or from multiple variant models, each representing a different combination of causal variants^2^. If the amount of statistical information at any trait-associated variant increases, e.g. by increasing sample size, the BF will tend to increase. Therefore, if there is variable information between variants in a sample, the BF are not directly comparable between variants. We show here that as GWAS sample sizes increase, very small differences in the data quality at different variants may cause large differences in the credible sets, even causing replacement of one variant in a single variant credible set by another. This tallies with results which showed that fine-mapping can be inaccurate applied to meta-analyses ^3^, where differential information may relate to different genotyping arrays and imputation quality.

One common approach to colocalisation, coloc ^4^, is based on fine-mapping two traits in a single region, and deriving posterior probabilities of colocalisation as functions of the fine mapping statistics. We show that that the sensitivity of fine-mapping to small differences in data quality propagates to coloc, and may cause incorrect, but apparently confident, inference of either colocalisation or non-colocalisation. We propose a solution to this problem, based on adapting a previous proposal for a proportional colocalisation test ^5,6^.

This test is sensitive to other issues of data quality, and is thus not a replacement for standard colocalisation analysis. But if both methods draw the same conclusion, this can provide greater confidence in inference, in order to design follow-up experiments.

Alternatively, if coloc produces high posterior support against colocalization, which is incompatible with other data and an investigator’s visual inspection of the data, support for colocalization by the proportional test may indicate a false coloc result due to differential information across variants.

## Results

### The effect of data quality on fine mapping and colocalisation inference

Variable information across variants can arise through differential genotyping error or completeness, but also through differential imputation quality. It can also arise from rounding summary statistics to a small number of significant figures, where the difference between the rounded and complete statistic then varies across variants. We conducted a simulation study where two variantss in complete linkage disequilibrium (LD) were simulated, and then a degree of information loss imposed on one of the variants. In the complete data, we would expect each variant to receive a posterior probability of 0.5.

However, we observed that the posterior probability deviated from the expected 0.5 as information loss increased, with the effect being stronger at higher sample sizes. For example, at an imputation r^2^ of 0.95, the posterior probability at the variant with information loss was about 0.45 in a 5,000 sample study, but had a median of 0 in a 500,000 sample study (Figure 1). This latter case would tend to produce a single variant credible set, with a 50% chance this would be incorrect. We see that increasing bias in posterior probability is associated with an increasing tendency to declare a single causal variant credible set (Supplementary Figure 1).

**Figure 1.**
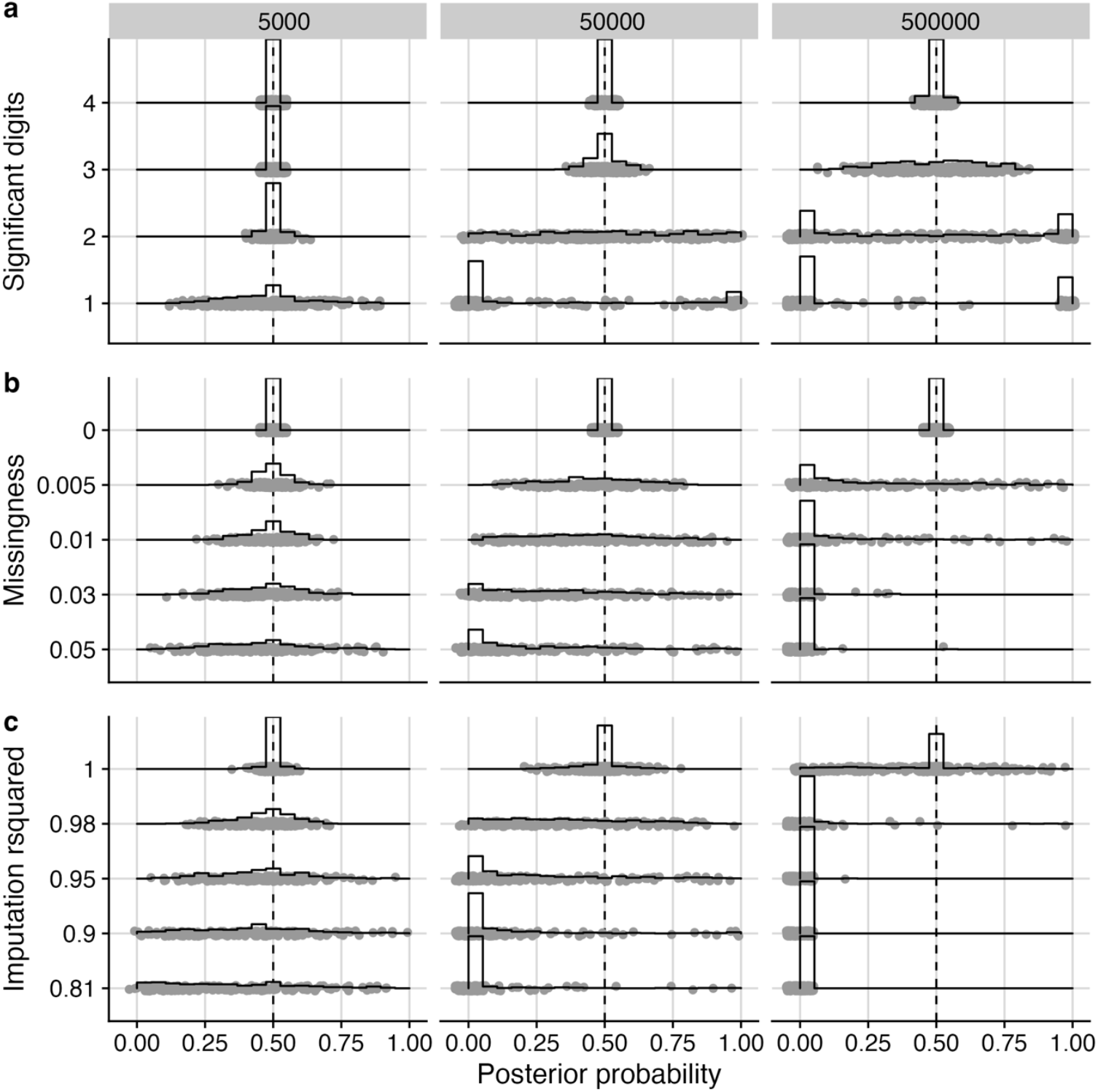
Effect of information loss on fine-mapping posterior probabilities. In all cases we simulated two variants in perfect LD, and imposed information loss on one of them, by reducing the number of significant digits stored for summary statistics, by reducing the fraction of genotyped samples, or by decreasing imputation r^2^. In each row, information decreases moving from top to bottom, and the distribution of posterior probabilities, expected to be 0.5 with complete information, is shown for the variant with information loss. Histograms are used to show the distribution of the posterior probabilities. The three columns differ by simulated sample size.

### Adaptation of a proportional test of colocalisation

A previously proposed proportional test of colocalisation^5,6^ assesses the null hypothesis that the effect sizes across any subset of variants in a region are proportional between the two traits. Thus, if the two traits give us effect estimates 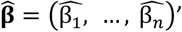 and 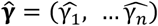 across the same set of *n* variantss, the null hypothesis is that

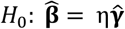

for some η. This is equivalent to saying we can draw a straight line, through the origin, through *n* points representing the true effects for each trait (Figure 2). The test for proportional colocalization is based on Fieller’s statistic

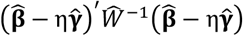

where 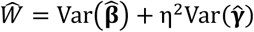 is the variance-covariance matrix of 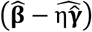. As η may be 0 or infinity, we use a change of variables to θ = tan^−1^ η, and use the test statistic

**Figure 2.**
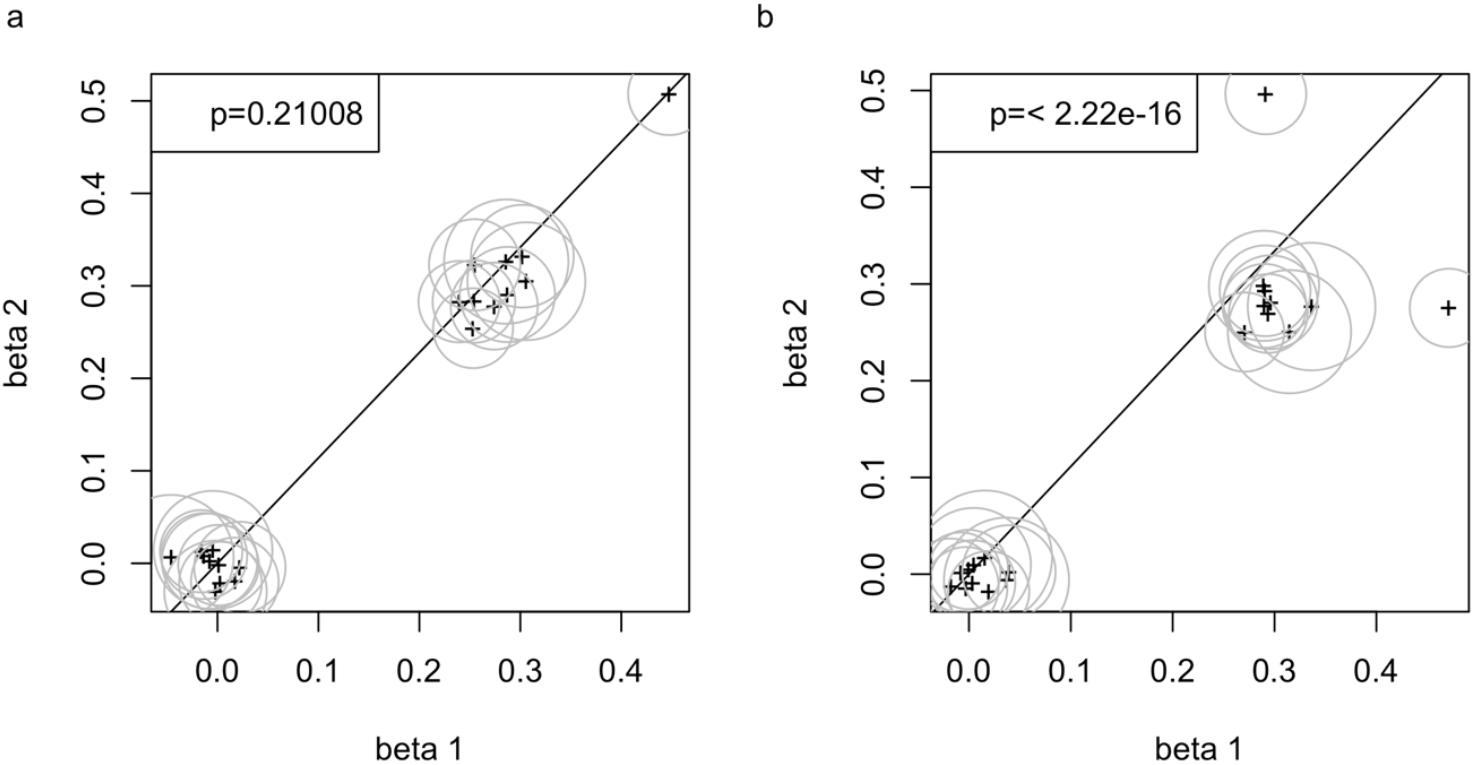
Examples of proportional colocalization testing. The x axis shows beta from study 1, the y axis the beta from study 2. Circles indicate 95% confidence regions. The line has slope 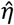. P values for the test of the null hypothesis of proportional colocalization are shown. **a** the causal variants are the same for the two studies (H0 is true). **b** the causal variants differ, but are in LD (H0 is false).

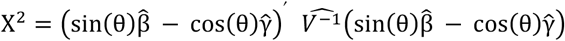

where 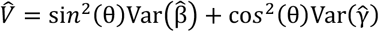. Replacing θ by its maximum likelihood estimate, 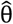, following a profile likelihood approach, means the test statistic has a 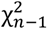 distribution. The advantage of this approach compared to a fine-mapping based approach is that variable information can be captured in the standard errors of the estimated effects and accounted for.

While the differential information is captured by the standard errors of the estimates, we need to pay special attention to patterns of genotype missingness, as these impact the covariance of 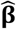 and 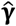, and hence 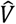. When there is no genotype missingness then 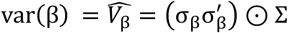 with Σ the LD matrix and σ the vector of standard errors of 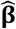 across variants being tested (and similar for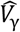). If two variants are missing a portion of their data, 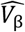 will differ depending on whether the missing data comes from the same samples for both variants, or different samples. We show in the supplementary text that we may scale the standard estimates of 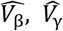 given above by a function of the number of shared samples across pairs of variants to get a more accurate estimate.

There are three challenges with this test which have limited the use of the proportional test in practice and which we address in this work. First, the original proposal assumed that 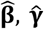, were estimates from a joint model of all *n* variants, requiring access to individual-level data. Here, we reframe the test to use the marginal test statistics and a reference LD matrix which means standard GWAS summary statistics can be used without need for reconstruction of the joint model statistics. Second, the null hypothesis is satisfied for the trivial cases where there is no genetic effect (β=0 or γ = 0), so we propose to only consider the proportional test when coloc has given strong evidence that there is association with both traits. Finally, the use of all *n* variants in a region will lead to a weakly powered test with *n-1* degrees of freedom. The original proposal^5^ used only two variants, the mostly strongly associated with each trait. Whilst this is indeed a powerful approach, it has been shown it also leads to very poor control of type 1 error rates^6^. Instead, a compromise was proposed where the *n* variantss are summarised by a smaller number of principal components^6^, to avoid the selection effect which induces bias. However, this is difficult to construct from marginal test statistics, and is still suboptimal in terms of degrees of freedom. Here, we propose to instead conduct all *n*(*n*-1)/2 possible tests of pairs of variants, and use a false discovery rate (FDR) to examine whether any of these are significant, rejecting the null hypothesis if it is rejected for any pair of variants. We show (Figure 3) that the p-value of the test is bounded below by the minimum GWAS test p-value for either trait across the variants to be tested. This enables us to only perform the test for a subset of variants, as long as we allow for the total number of possible tests in the FDR procedure.

**Figure 3.**
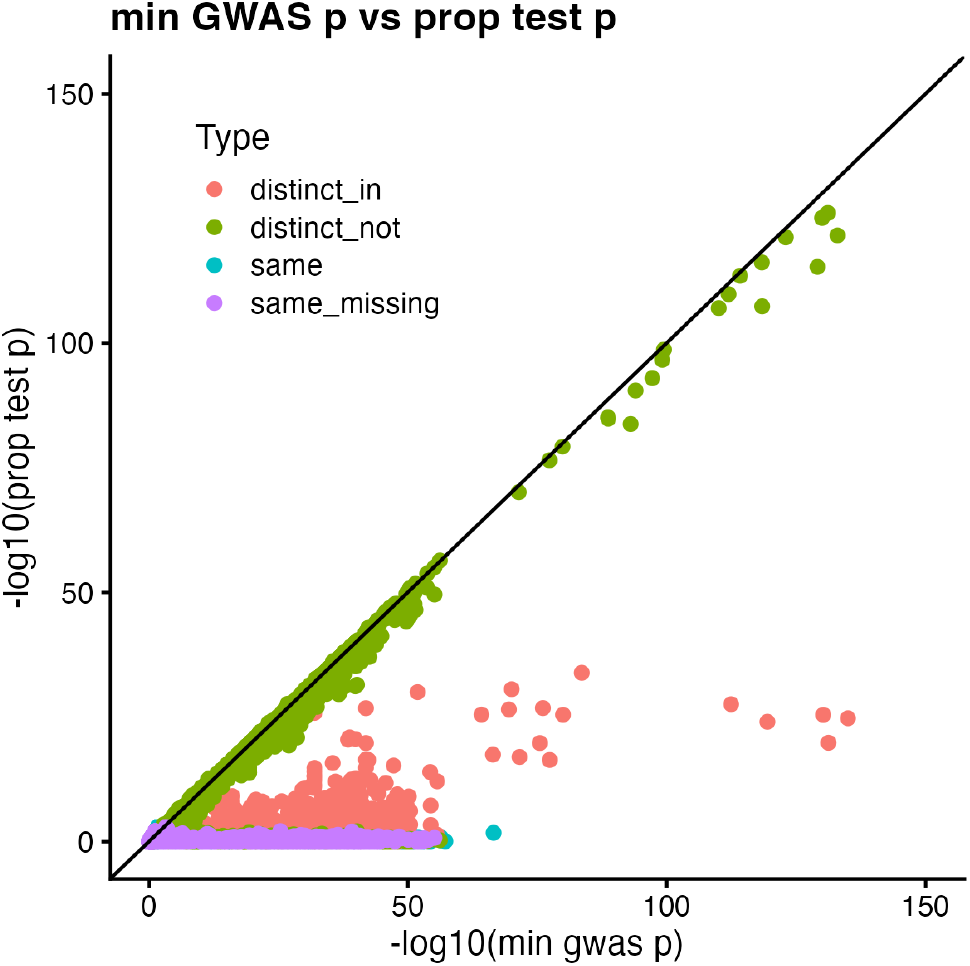
The proportional test p-value is bounded below by the minimum marginal p-value across variants used in a proportional test. The plot compares p-values from colocPropTest (y axis) with the minimum p-value over the pair of SNPs and pair of traits contributing to colocPropTest (x axis). The different colours relate to simulation “Type” - whether the causal variant in two simulated datasets was distinct but in some LD (distinct_in), distinct but not in LD (distinct_not), the same (same) or the same with differential genotype missingness which could bias fine-mapping based coloc analysis (same_missing).

### Simulations show the proportional test can correctly resolve false coloc inference

We simulated GWAS summary data for two traits for small regions under three scenarios: that the causal variants were the same (A), distinct but in LD (B), or distinct and not in LD (C). Where the causal variants were the same, we imposed missing genotype coverage on one dataset at the most strongly associated variants – either independently across the variants (A1) or in the same subset of samples (A2). We found that coloc made correct inference when causal variants were distinct and not in LD (Figure 4). When they were distinct but in LD, we would expect coloc to call H3, but there was a risk of up to 25% of falsely calling H4 as genotype missingness as LD increased. When they were the same, and we would expect to call H4, there was a risk up to 100% of calling H3, as genotype missingness and LD increased. On the other hand, the proportional test rejected the null of colocalisation consistently when causal variants were distinct (B, C). The proportional test typically accepted the null of colocalisation when causal variants were the same (A1, A2), although there was a loss of power at high levels of missingness in the presence of high LD. The pattern was similar regardless of how that missingness was imposed (using the same or different samples across the variants).

**Figure 4.**
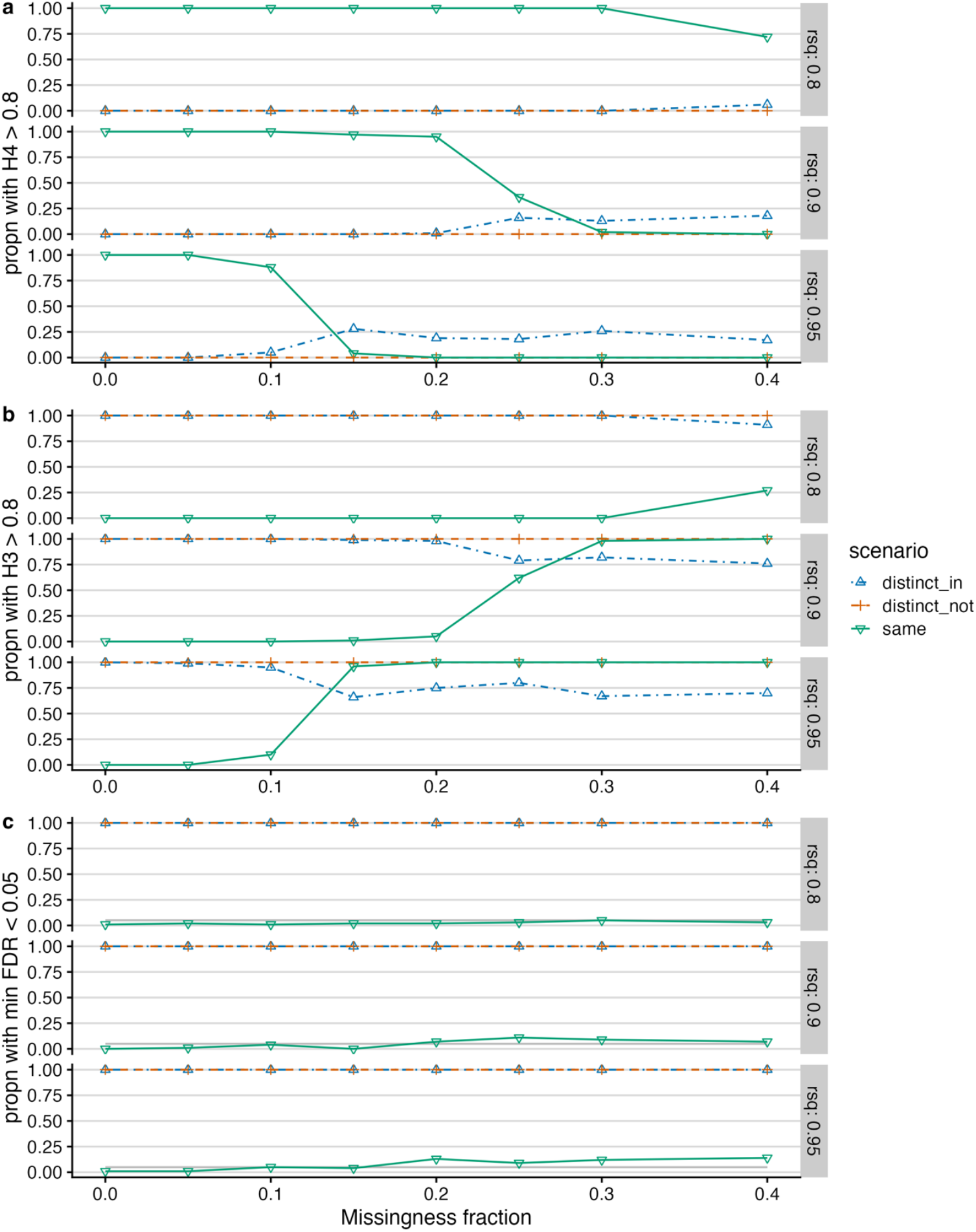
Comparison of coloc and proportional colocalisation tests in simulated scenarios. We considered two datasets each with a single causal variant. These causal variants might be distinct between the datasets and either in LD (distinct_in) or not in LD (distinct_not), or the same. Each simulation consisted of blocks of variants in with r^2^ 0.8, 0.9 or 0.95 (rows). Tests were performed on data with genotype missingness imposed independently on the most significant variants in the second dataset. Plots a and b show the coloc results in terms of posterior probabilities for the H4 (shared) and H3 (distinct) hypotheses, while plot c shows the proportion of simulations for which the proportional colocalisation FDR was below 0.05. We simulated 100 examples of each scenario. See Supplementary Figure X for the same analysis with missingness imposed on the same samples across variants in the second dataset.

## Discussion

Fine-mapping and colocalization methods were originally developed when GWAS sample sizes were at least an order of magnitude smaller than today. In smaller sample sizes, issues with variable information across SNPs might produce slight bias in fine-mapping probabilities, but as credible sets were generally large, the causal variant was likely to lie within the credible set. However, in modern GWAS which may be formed from meta-analysis across multiple biobank studies as well as trait-specific studies, credible sets of single variants have become more common. These offer apparent confidence in identifying causal variants that may not be warranted. The errors propagate to coloc which may provide apparently confident inference (posterior probabilities close to 1) that are incorrect.

The proportional test allows an alternative approach to colocalisation that overcomes the issue of variable information by incorporating a measure of that information, the standard error, as a separate parameter in its test statistic. Our adaptation here to maintain power is the use of multiple pairwise tests paired with a false discovery rate approach. An alternative extension of the proportional colocalisation test was recently published which allows selection of specific variants (e.g. by minimum single trait p-values), and then adjusts for the selection to reduce the previously reported bias^7^.

Proportional tests will be sensitive to departures from different assumptions than fine-mapping-based colocalization. For example, coloc does not assume that both GWAS data have equal LD, whereas proportional testing will require equal LD. Because we reject the null if any one test statistic is large enough, we must ensure all data entering the proportional test is good quality, so we advocate that strong quality control is applied independently to each dataset. Screening by DENTIST^8^ has been shown to increase accuracy of fine-mapping.

Careful attention must also be paid to the allele matching of SNPs between datasets: for example if a triallelic SNP has been coded differently between the datasets, this could cause false rejection of the null. On the other hand, unlike approaches based upon fine-mapping, the test does not assume that the true causal variant is contained in the dataset, so density of variant coverage is less of an issue (if the true causal variant is not contained in the dataset, we assume some variant(s) in LD with it are – if the true causal variant is not in the dataset and not in LD with any variants in the dataset, neither colocalisation approach has information to make accurate inference).

However, proportional methods work by avoiding any fine-mapping step. New methods are needed to enable robust fine-mapping of large datasets where very small variability in information between variants can produce very confident, incorrect inference.

## Methods

### Fine-mapping simulations

We simulated genotypes for a single binomial variable with minor allele frequency uniformly sampled in the interval (0.05,0.5) and assigned that value to each of the two variant variables. We simulated a case-control outcome variable as a Bernoulli variable with a 20% frequency in the common homozygote, and an odds ratio sampled from a normal distribution with mean 1.2 and standard deviation 0.02. We calculated summary statistics at each variant, and from these an approximated Bayes factor using Wakefield’s method^1^. We calculated posterior probabilities at each variant as the Bayes factor for that variant divided by the sum of the Byes factors across the variants. We imposed missing information on the second copy of the genotype in one of three ways:

1. Rounding the summary statistics to 1, 2, 3 or 4 decimal places
2. Removing genotypes from a fraction of samples (missingness)
3. Replacing the genotype with a vector imperfectly correlated with it, where we recorded the r^2^ between the original genotype and the imperfect copy.

We compared sample sizes of 5,000, 50,000 and 500,000.

### Bounded p-value simulations

We simulated individual level GWAS data for two traits under a range of scenarios, and conducted colocPropTests for every pair of variants. We compared the resulting test p-value against the “maximin p value’’, defined as the maximum over traits of the minimum p-value over variants for each trait.

### Colocalisation simulations

We simulated genotypes for 5000 individuals at ten variants in strong LD, and an independent set of another ten variants in strong LD – i.e. each region consisted of 20 vairants. One variant was randomly chosen as the causal variant for the first trait. For the second trait choice of causal variant varied with the scenario:

- A: the variant was chosen to be the same
- B: to be distinct in the same block (“distinct_in”)
- C: distinct in the other block (“distinct_not”)

Genotypes were simulated as correlated binomial samples, generated as the sum of two sets of correlated Bernoulli variables. A normally distributed quantitative trait was simulated and summary statistics calculated.

When the scenario was the for the same causal variant in each trait with missing data, we imposed variable information across variants for the second trait as follows. We identified the three most significant variants, and removed genotype data for 80% of individuals, either

- A1: randomly across the three variants (“same_miss”) or
- A2: in a coordinated manner, selecting the same 80% of individuals to have data hidden across the three variants (“same_miss2”)

This extreme level of missingness was chosen to ensure coloc would consistently miscall “same_miss” and “same_miss2” as H3 (distinct causal variants). We applied standard coloc with default settings, and the proportional test set out above using the colocPropTest package.

## Availability of Code

The colocPropTest package is available in CRAN: https://cran.r-project.org/web/packages/colocPropTest/index.html. Full code for simulations is available at https://github.com/chr1swallace/propcoloc-manuscript/.

## Conflicts of Interest

All authors are employees of GSK.

