## Supplementary Information for "Variable information across SNPs in GWAS data can cause false rejections of colocalisation which can be resolved by proportional colocalisation tests"

June 16, 2025

**2 Supplementary Figures** **6**

### 1 Description of test statistic

Given a vector of regression coefficients, $\hat{\beta}$, and its standard error $\sigma_\beta$, at a set of SNPs, and the same for another trait $\hat{\gamma}, \sigma_\gamma$, Fieller's test statistic, which assumes $\hat{\beta} \perp\!\!\!\perp \hat{\gamma}$ is

$$(\hat{\beta} - \eta\hat{\gamma})' W^{-1} (\hat{\beta} - \eta\hat{\gamma})$$

where

$$W = \operatorname{var}(\hat{\beta}) + \eta^2 \operatorname{var}(\hat{\gamma})$$

To avoid discontinuities where $\eta$ is 0 or $\infty$, we employ a change of variables, $\theta = \tan^{-1}(\eta)$. Then the test statistic becomes

$$(\sin(\theta)\hat{\beta} - \cos(\theta)\hat{\gamma})' V^{-1} (\sin(\theta)\hat{\beta} - \cos(\theta)\hat{\gamma})$$

with

$$V = \sin^2(\theta) \operatorname{var}(\hat{\beta}) + \cos^2(\theta) \operatorname{var}(\hat{\gamma})$$

We can estimate $V = V_\beta + V_\gamma$ where

$$V_\beta = (\sigma_\beta \sigma_\beta') \odot \Sigma$$

where $\Sigma$ is the LD matrix of snp-snp correlations, and similarly for $V_\gamma$.

### 1.1 Overlapping samples

If $\hat{\beta} \not\perp \hat{\gamma}$, then

$$V = \sin^2(\theta)\,\mathrm{var}(\hat{\beta}) - \cos^2(\theta)\,\mathrm{var}(\hat{\gamma}) - \sin(\theta)\cos(\theta)\,\mathrm{cov}(\hat{\beta}, \hat{\gamma})$$

and we can estimate

$$\mathrm{cov}(\hat{\beta}, \hat{\gamma}) = \rho(\sigma_\beta \sigma'_\gamma) \odot \Sigma$$

where $\rho$ is the correlation between $Z$ scores between the two datasets, with the correlation taken over a large set of SNPs.

### 1.2 Effect of variable genotype missingnes on correlation between effect estimates at a pair of SNPs

If two SNPs are in LD, with correlation coefficient $\rho$, conveniently, the Z scores from regressing any phenotype onto each SNP in turn will also have expected correlation $\rho$.

However, this assumes the two SNPs are genotyped in the same samples, which may not happen when meta analysis is involved. We evaluated this using simulation. We simulated genotypes at two SNPs as correlated binomial variables, $X_1, X_2$ and an outcome variable $Y$ either dependent on $X_1$ or independent of both $X$. We computed regression coefficients for each complete data example, then hid a fraction of genotypes and recomputed them. We repeated this 100 times for different combinations of minor allele frequency (MAF), LD, and missingness fraction (Table 1) to calculate the correlation between regression coefficients in the reduced dataset. The simulations showed that when a different subset of samples are genotyped for two SNPs, the reference dataset $r$ could overestimate the empirical correlation (Figure 1).

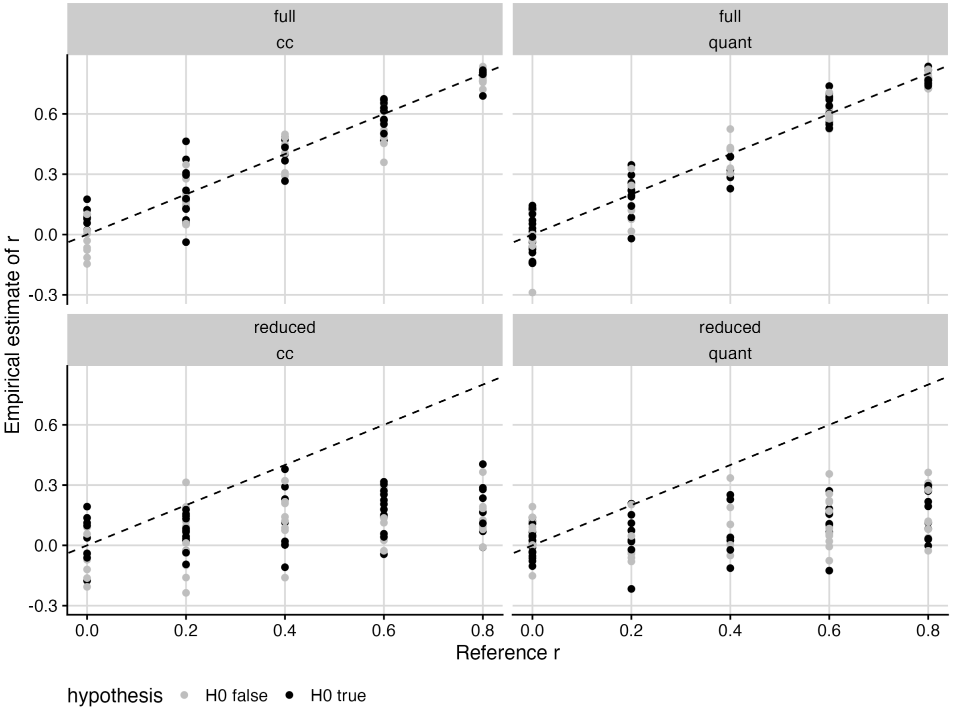

Figure 1: Comparison between population LD ($r$) and empirical estimates of correlation of test statistics in full and downsampled data

Here, we estimate how partial genotype missingness may affect the correlation between regression coefficients, following workings from [Lin and Sullivan, 2009].

Assume we have snps $k$, $l$, with the number of cases and controls genotyped at SNP $k$ given by $N_k^1, N_k^0$ respectively (and similarly for snp $l$). We use the notation $N_{lk}^*$ for the number of samples in common between snps $l$ and $k$.Following Lin and Sullivan, let $\theta_k, \theta_l$ be the regression coefficients, estimated from a vector of cases $Y$ and genotypes $X$. To allow an intercept term, we augment the $X$ by prepending a vector of 1, to give $\tilde{X}$.

At SNP $k$ the logistic likelihood is

$$L(\theta_k) = \prod_i \frac{e^{Y_i(\alpha_k+\beta_k X_{ik})}}{1+e^{\alpha_k+\beta_k X_{ik}}}$$

with score function

$$U(\theta) = \sum_i \left(Y_i - \frac{e^{\alpha_k+\beta_k X_{ik}}}{1+e^{\alpha_k+\beta_k X_{ik}}}\right)\begin{pmatrix}1\\X_{ik}\end{pmatrix}$$

and information matrix

$$I(\theta_k) = \sum_i \frac{e^{\alpha_k+\beta_k X_{ik}}}{(1+e^{\alpha_k+\beta_k X_{ik}})^2}\begin{pmatrix}1 & X_{ik}\\X_{ik} & X_{ik}^2\end{pmatrix}$$

and similarly for SNP $l$. Then,

$$\text{cov}(\theta_l,\theta_k) = I_k^{-1}(\theta_k)cov(U_k(\theta_k),U_l(\theta_l))I_l^{-1}(\theta_l)$$

We assume that $H_0$ is true (i.e. $\beta_k=\beta_l=0$). We write $f_k, f_l$ for the MAF of snps $k,l$. Note that the estimated MAF will vary with the sample, but we make the simplifying assumption that all estimated MAF are approximately equal for a given SNP. This follows from the assumption that we are operating under $H_0$, ie that there are no systematic differences between allele frequency in cases and controls.

#### 1.2.1 Estimation

Under $H_0$,

$$I_k(\theta_k) = \frac{N_k e^{\alpha_k}}{(1+e^{\alpha_k})^2}\begin{pmatrix}1 & 2f_k\\2f_k & 2f_k(1+f_k)\end{pmatrix} = \frac{N_k e^{\alpha_k}}{(1+e^{\alpha_k})^2}H_k,$$

where $H_k$ is a matrix only dependent on allele frequencies at SNP $k$, so that

$$I_k^{-1}(\theta_k) = \frac{(1+e^{\alpha_k})^2}{N_k e^{\alpha_k}}H_k^{-1}$$

and similarly for $I_l^{-1}(\theta_l)$.

Also

$$\begin{aligned}\text{cov}(U_k(\theta_k),U_l(\theta_l)) &= \sum_{i=1}^{N_{lk}}\left(Y_i - \frac{e^{\alpha_k}}{1+e^{\alpha_k}}\right)\left(Y_i - \frac{e^{\alpha_l}}{1+e^{\alpha_l}}\right)\tilde{X}_{ki}\tilde{X}_{li}^T\\
&= \frac{e^{\alpha_k+\alpha_l}}{(1+e^{\alpha_k})(1+e^{\alpha_l})}\begin{pmatrix}N_{lk} & \sum X_{li}\\ \sum X_{ki} & \sum X_{li}X_{ki}\end{pmatrix} + \qquad \textit{sum over all shared samples}\\
&\left(1 - \frac{e^{\alpha_l}}{1+e^{\alpha_l}} - \frac{e^{\alpha_k}}{1+e^{\alpha_k}}\right)\begin{pmatrix}N_{lk}^1 & \sum X_{li}\\ \sum X_{ki} & \sum X_{li}X_{ki}\end{pmatrix} \qquad \textit{sum over all shared cases}\\
&= \frac{N_{kl}e^{\alpha_k+\alpha_l} + N_{kl}^1(1-e^{\alpha_k+\alpha_l})}{(1+e^{\alpha_l})(1+e^{\alpha_k})}H\end{aligned}$$

| Parameter | possible values |
|---|---|
| MAF | 0.1, 0.2, 0.3, 0.4, 0.5 |
| LD ($r$) | 0, 0.2, 0.4, 0.6, 0.8 |
| missingness fraction | 0.6, 0.7, 0.8, 0.9 |

Table 1: Values of simulation parameters

where $H$ is a matrix only dependent on allele frequencies which we assume are independent of sample choice under $H_0$. Thus,

$$\begin{aligned}\mathrm{cov}(\theta_k, \theta_l) &= \frac{(1+e^{\alpha_k})^2(1+e^{\alpha_l})^2}{N_k e^{\alpha_k} N_l e^{\alpha_l}} \frac{N_{kl} e^{\alpha_k+\alpha_l} + N_{kl}^1(1-e^{\alpha_k+\alpha_l})}{(1+e^{\alpha_k})(1+e^{\alpha_l})} H_k^{-1} H H_l^{-1} \\ &= \frac{(1+e^{\alpha_k})(1+e^{\alpha_l})}{N_k N_l e^{\alpha_k+\alpha_l}} (N_{kl}^1 + N_{kl}^0 e^{\alpha_k+\alpha_l}) H_k^{-1} H H_l^{-1}\end{aligned}$$

Now $V(\theta_k) = I_k^{-1}(\theta_k)$ (and similar for $V(\theta_l)$) so

$$\begin{aligned}\mathrm{cor}(\theta_k, \theta_l) &= \frac{(1+e^{\alpha_k})(1+e^{\alpha_l})}{N_k N_l e^{\alpha_k+\alpha_l}} (N_{kl}^1 + N_{kl}^0 e^{\alpha_k+\alpha_l}) \frac{\sqrt{N_k N_l e^{\alpha_k} e^{\alpha_l}}}{(1+e^{\alpha_k})(1+e^{\alpha_l})} H_k^{-1} H H_l^{-1} \left(H_k^{-1} H_l^{-1}\right)^{-1/2} \\ &\simeq \frac{N_{kl}^1 + N_{kl}^0 \frac{N_k^1 N_l^1}{N_k^0 N_l^0}}{\sqrt{N_k N_l \frac{N_k^1 N_l^1}{N_k^0 N_l^0}}} P \\ &= \frac{N_{kl}^1 \sqrt{\frac{N_k^0 N_l^0}{N_k^1 N_l^1}} + N_{kl}^0 \sqrt{\frac{N_k^1 N_l^1}{N_k^0 N_l^0}}}{\sqrt{N_k N_l}} P\end{aligned}$$

where we use $e^{\alpha_k} \simeq N_k^1 / N_k^0$ and similar for $e^{\alpha_l}$, and write $P = H_k^{-1} H H_l^{-1} \left(H_k^{-1} H_l^{-1}\right)^{-1/2}$.

For quantitative traits, a similar but simpler derivation gives

$$\mathrm{cor}(\theta_k, \theta_l) = \frac{N_k l}{\sqrt{N_k N_l}} P'$$

We can use these estimaters to adjust a given reference value of $r$, by dividing by $cor(\theta_k, \theta_l)$ given full sample size and multiplying by the same calculated given observed sample size (note that $P$ and $P'$ cancel). Our challenge is to calculate $N_{kl}^*$, the numbers of cases and controls typed for both SNPs. We approximate this by $\min(N_k^*, N_l^*)$ in the absence of other information.

#### 1.2.2 Evaluation

We used the same simulations as above, this type predicting between estimate correlation using the calculations above. We found agreement was good, validating our calculations (Figure 2).

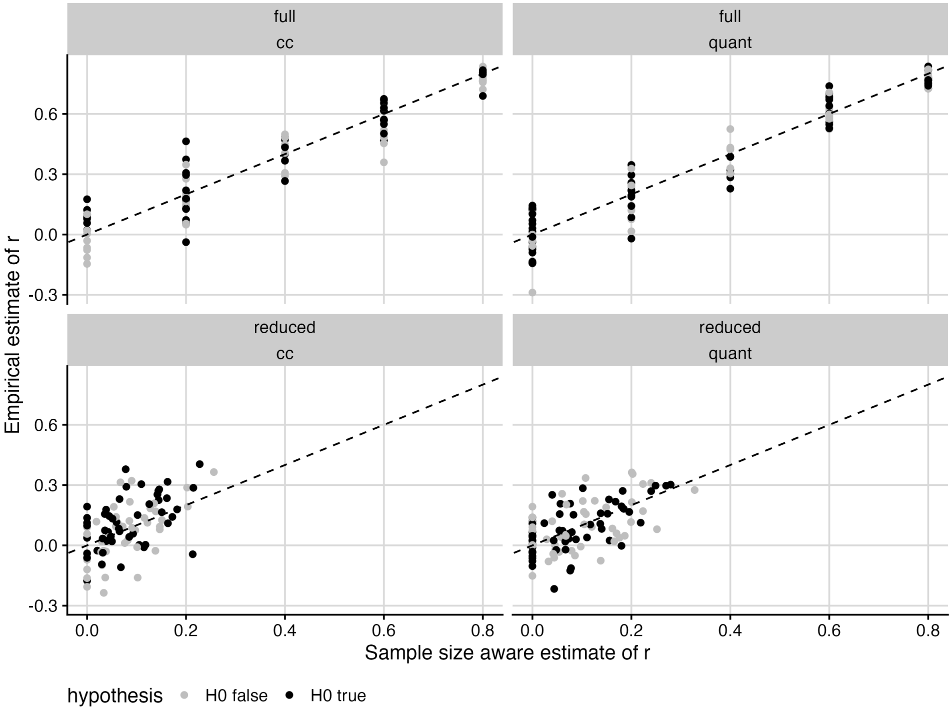

Figure 2: Comparison between population LD ($r$) and theoretical estimates of correlation of test statistics in full and downsampled data using sample-size aware estimates above

### 2 Supplementary Figures

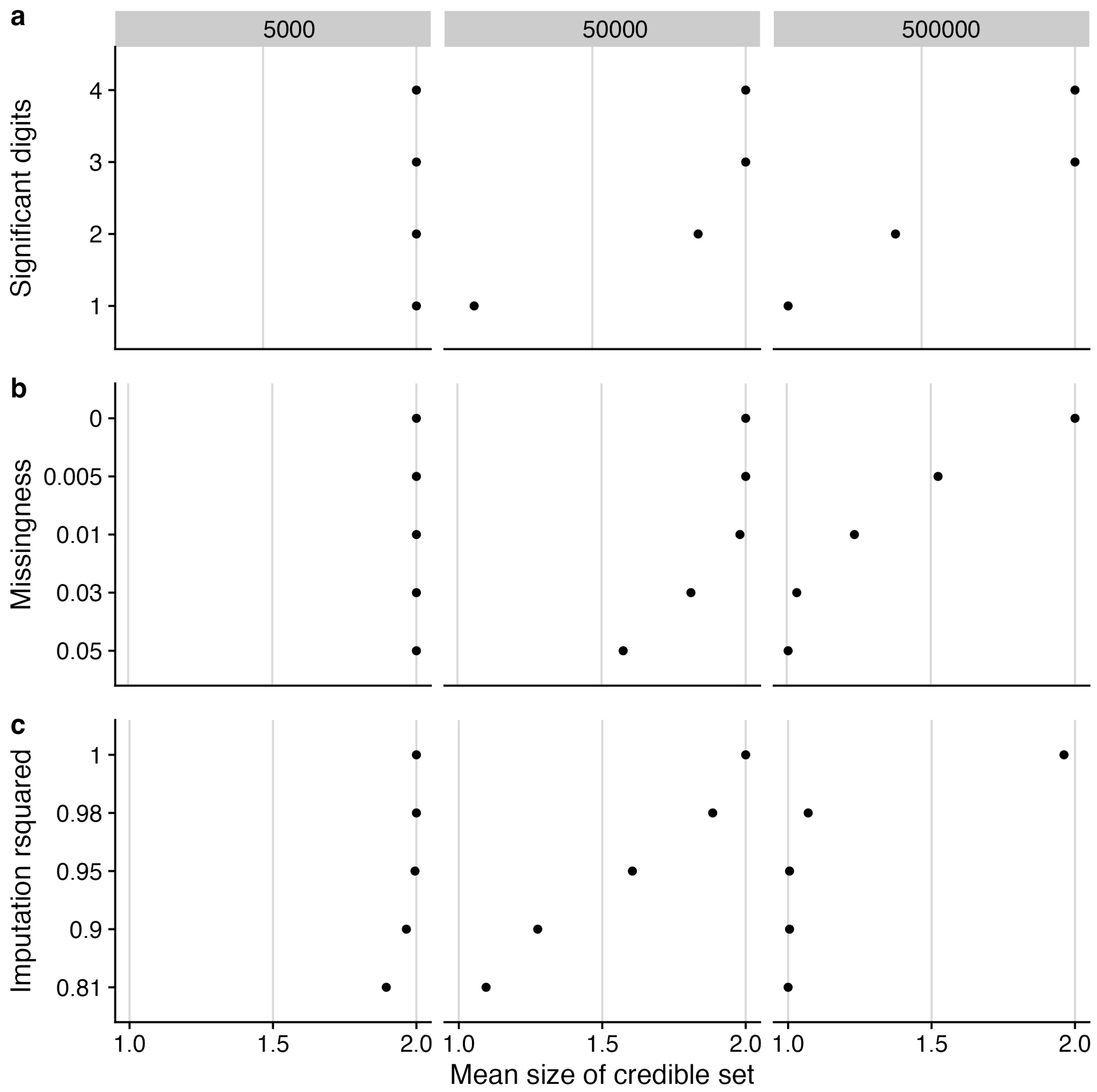

Figure 3: Effect of information loss on credible set size. Same data as in Figure 1. In all cases, we expect the credible set to contain 2 SNPs with complete data, and points show the mean number of SNPs in a 95% credible set over 200 simulated datasets.

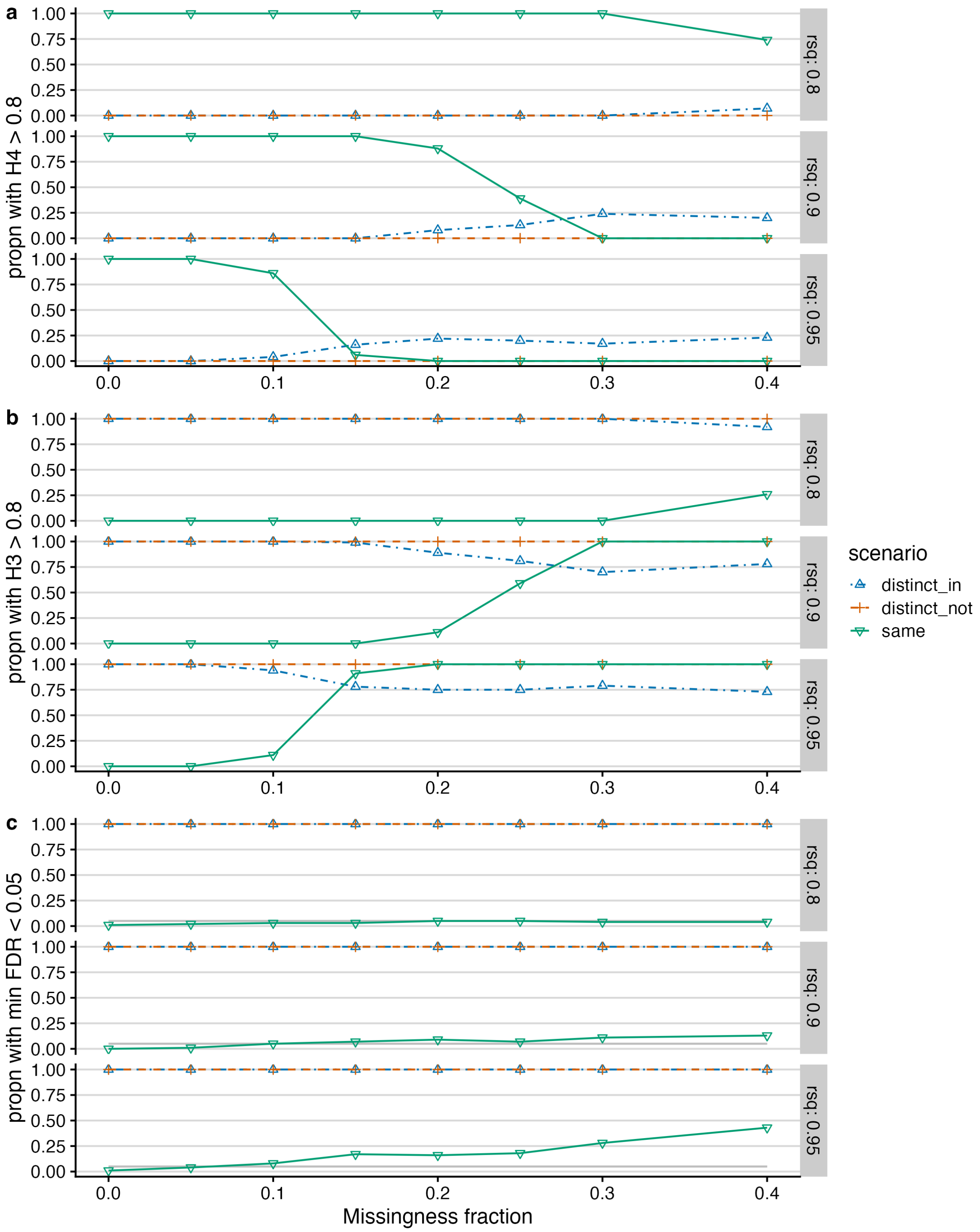

Figure 4: Companion to Figure 4, showing results when missing data is imposed randomly across samples in the second dataset instead of across the same samples.
